# Improved Nanopore full-length cDNA sequencing by PCR-suppression

**DOI:** 10.1101/2022.07.28.501892

**Authors:** Anthony Bayega, Spyros Oikonomopoulos, Yu Chang Wang, Jiannis Ragoussis

## Abstract

Full-length transcript sequencing remains a main goal of RNA sequencing. However, even the application of long-read sequencing technologies such as Oxford Nanopore Technologies still fail to yield full-length transcript sequencing for a significant portion of sequenced reads. Since these technologies can sequence reads that are far longer than the longest known processed transcripts, the lack of efficiency to obtain full-length transcripts from good quality RNAs stems from library preparation inefficiency rather than the presence of degraded RNA molecules. It has previously been shown that addition of inverted terminal repeats in cDNA during reverse transcription followed by single-primer PCR creates a PCR suppression effect that prevents amplification of short molecules thus enriching the library for longer transcripts. We adapted this method for Nanopore cDNA library preparation and show that not only is PCR efficiency increased but gene body coverage is dramatically improved. The results show that implementation of this simple strategy will result in better quality full-length RNA sequencing data and make full-length transcript sequencing possible for most of sequenced reads.

**Contribution to the field:** Long-read RNA sequencing aims to sequence expressed transcripts in their entirety. However, this has remained a challenge, mainly due to inherent inefficiencies in cDNA library preparation. Herein, we provide a new Nanopore cDNA library preparation protocol, termed Panhandle, that improves the efficiency of cDNA PCR with yields 2 – 8 times the yields obtained with ordinary PCR. This is key, as this should in turn reflect in the possibility of lowering the number of PCR cycles needed to obtain ample sequencing material, which in turn could reduce PCR biases, PCR artifacts, turnaround time, reagents, and could increase general quality of the library. Further, transcripts generated using the Panhandle method show better gene body coverage and more accurate transcription start site mapping than regular methods. This represents an important step towards full-length cDNA sequencing by Nanopore.

## Introduction

Since its discovery in 1961 (Brenner et al., 1961), RNA remains the subject of intensive research, while finding application in therapeutic developments, as well as a tool for clinical diagnostics development. RNA sequencing (RNAseq) is the most complete way to analyze gene expression by determining absolute and relative abundances of transcripts, as well as by identifying isoforms (Stark et al., 2019). Majority of RNAseq experiments are performed using short-read sequencing technologies. However, since the lengths of most transcripts surpass the length attainable by currently available short-read sequencing technologies, long-read sequencing technologies, such as Oxford Nanopore Technologies (ONT) have been employed (Bayega, Fahiminiya, et al., 2018; Oikonomopoulos et al., 2020). For example, majority of human transcripts are 1 – 2 kb in length with the longest known processed human transcript, the Titin mRNA, stretching over 100 kb (Bang et al., 2001). The current recorded longest read sequenced with Oxford Nanopore Technologies (ONT) platform is over 2 Mb (Payne et al., 2018), suggesting that long-read RNAseq should yield full-length transcripts. However, as currently applied, long-read sequencing technologies fail to yield full-length cDNA sequences for up to 50 % of sequenced reads (Chen et al., 2021). Since the read lengths attainable by long-read sequencing technologies surpass the lengths of most transcripts, the bottleneck in obtaining full-length transcripts for most sequenced reads, provided very good original RNA quality, appears to be technical inability to prepare long enough transcripts for sequencing.

In its most classical application, RNAseq involves the isolation and purification of total RNA followed by conversion of RNA to cDNA using a reverse transcriptase. The cDNA is then amplified through polymerase chain reaction (PCR) followed by sequencing of the purified amplicons (Bayega, Wang, et al., 2018). Although advanced methods now exist that allow direct RNA (Garalde et al., 2018) and cDNA sequencing (Chen et al., 2021), the overwhelming majority of RNAseq experiments use PCR-amplified cDNA. One well known drawback of PCR when amplifying complex DNA mixtures, such as cDNA libraries, is the tendency to preferentially amplify short fragments at the cost of long ones thus biasing the representation of the cDNA library towards shorter molecules (Shagin et al., 1999). In our hands, we have observed overrepresentation of short fragments in amplified cDNA, which suggests a PCR bias. In order to overcome the effect of this bias on sequencing, some protocols, like the one used for Pacific Biosciences’ (PacBio) Iso-seq, have advised size selection of the library into partitions of pre-selected size ranges (Gordon et al., 2015) or in combination with 5′ cap selection (Cartolano et al., 2016). This is, however, disadvantageous as it biases species representation, adds extra laborious and expensive steps, and potentially might lead to RNA degradation during extra processing.

In order to improve representation of long fragments in complex mixtures other approaches have been tried. For example, it was observed that during single primer PCR of heterogeneous cDNA libraries, self-annealing structures are formed that decrease PCR efficiency. Lukyanov et al., (Lukyanov et al., 1995) took advantage of this phenomenon to add inverted terminal repeats (ITR) to the ends of cDNA. During the annealing phase of each PCR cycle, the 5′ and 3′ ends of single stranded ITR-modified molecules self-anneal, forming panhandle structures (Figure 1). The stability of these panhandle structures is dependent on the length of the molecule such that shorter molecules form more stable panhandle structures than longer molecules. The more stable panhandle structures formed by shorter molecules prevent primer binding thus reducing the amplification efficiency of short molecules in what is referred to as PCR suppression effect (Lukyanov et al., 1995). It was further reported that ITRs did not reduce the efficiency of amplification of individual sequences present in the initial RNA sample at different abundancies (Lukyanov et al., 1995). Varying the GC content of the ITR and primer and varying primer concentration, Shagin et al (Shagin et al., 1999) showed that the degree of PCR suppression effect could be regulated and thus, one could vary the average length of complex mixtures of DNA. In the current work, we adapt this method to Nanopore cDNA library preparation and show that, indeed, this approach improves full-length cDNA sequencing by Nanopore compared to conventional standard and widely used Nanopore community methods (Chen et al., 2021). We incorporate ITR sequences to our cDNA molecules during reverse transcription and use single-primer PCR for cDNA amplification, followed by Nanopore library preparation. We compare this method, which we refer to as Panhandle (or just Panh), to Oxford Nanopore Technologies’ SQK-PCB109 method, which we refer to as ONT. The Panhandle method showed more than 2-fold increase in cDNA amplicon yield (suggesting improved PCR efficiency) and led to improved gene body coverage.

**Figure 1:**
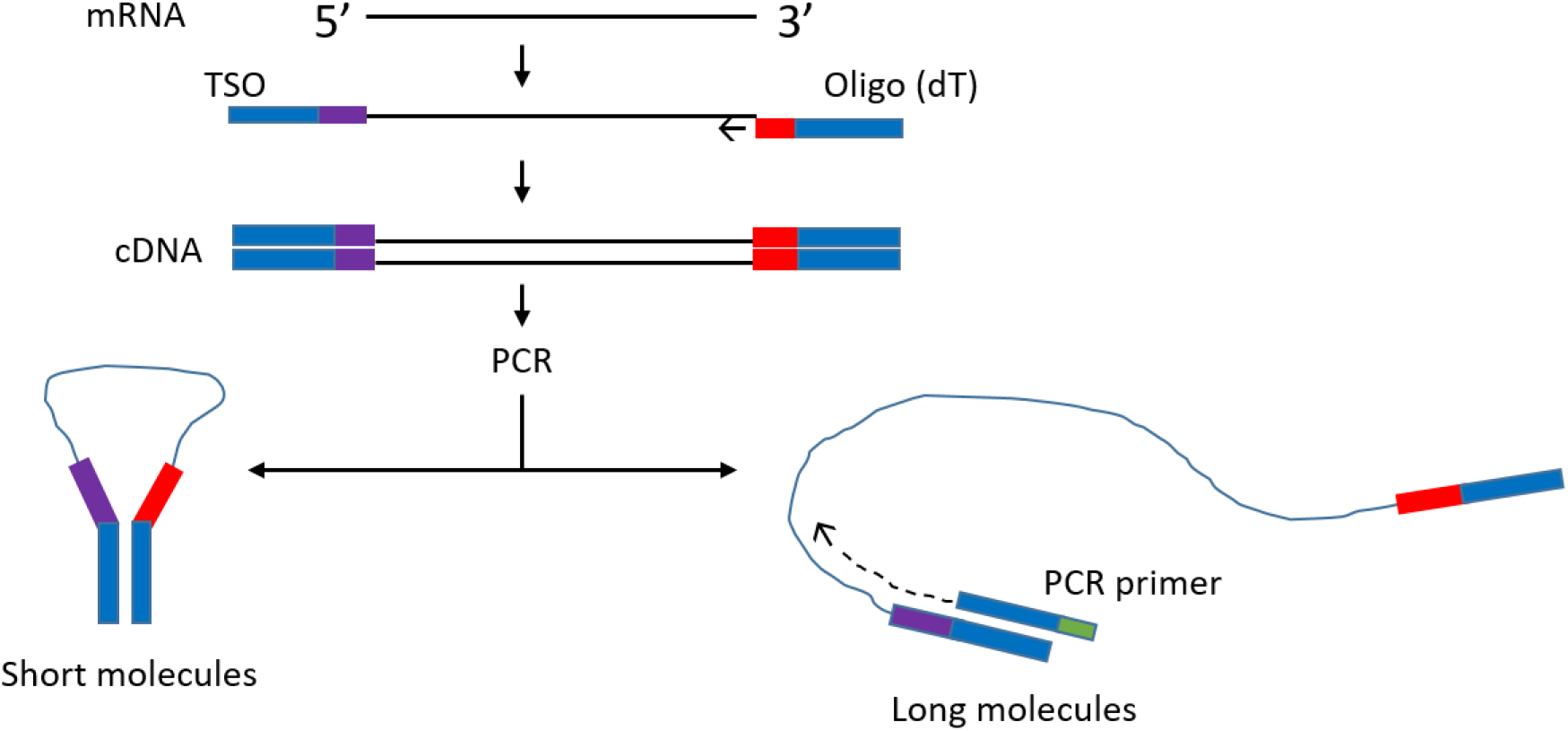
Schematic illustration of the Panhandle structure formation. During cDNA synthesis, we incorporate inverted terminal repeats in the oligo (dT) primer and template switching oligo (TSO) as shown with blue bars (green shaded region in **Table 1**). A single primer is used in PCR with complementary regions to the ITR present at both 5′ and 3′ ends of cDNA. Short cDNA fragments largely fail to denature preventing primers from binding. Long fragments denature allowing primers to bind as detailed by (Shagin et al., 1999). Adapted form (Shagin et al., 1999).

## Materials and methods

We obtained total RNA from four samples: Mediterranean fruit fly (Medfly) embryos, the human breast adenocarcinoma cell line MCF7 (Biochain, Newark, CA), HeLa cell line (SuperScript™ IV kit, Thermo Fisher Scientific), and a genome in a bottle sample, GM24143 (NIST, USA). Medfly total RNA was extracted from a pool of embryos collected at 6 hours post oviposition as previously described (Bayega et al., 2021). Total RNA was extracted from the female Epstein-Barr virus transformed B-lymphocyte cell line GM24143 using the Chemagic™ RNA TissuelO Kit H96 (PerkinElmer) following manufacturer’s instructions. MCF7 purified total RNA was purchased from Biochain (Newark, CA) while HeLa RNA was obtain from the Superscript IV kit (Thermo Fisher Scientific). We used 100 ng of Medfly total RNA, 50 ng of HeLa, 83 ng of GM24143, and 50 ng of Medfly total RNA.

In the ONT experiment, we processed the four samples following the Oxford Nanopore SQK-PCB109 kit according to manufacturer instructions. The samples were barcoded according to protocol, pooled at equal concentration, and sequenced on a single pre-used and washed PromethION flow cell which had 2521 pores. In the Panhandle experiment, we followed our in-house protocol as previously described (Bayega et al., 2021). The full protocol is added in full as Supplementary Protocol. Briefly, for each sample total RNA was added together with 1 μl of 10 μM oligo(dT) primer and 1 μl of 10 mM dNTPs in a 11.6 μl pre-RT reaction. The reaction was incubated at 72 °C for 3 minutes followed by 4 °C for 10 minutes, 25 °C for 1 minute and then held at 4 °C. A 10.4 μl reverse transcription (RT) reaction containing 1 X Maxima H Buffer, 1 μl RNaseOut (NEB), 2 μl of 100 μM TSO, 2 μl of 5M Betaine (Sigma-Aldrich), and 1 μl of Maxima H reverse transcriptase was added to the pre-RT reaction and the reaction incubated as shown in Supplementary Protocol. Following reverse transcription, 5 μl of cDNA was used in a 50 μl PCR reaction containing 1 μl of 10 μM PCR primer and 25 μl of 2x LongAmp Taq Master mix (NEB). PCR was performed as shown in Supplementary Protocol. The primers used in the Panhandle protocol are shown in Table 1. In both ONT and Panhandle approaches 20 PCR cycles were used. Following PCR, 1 μl of exonuclease (NEB) was added to each reaction and incubated for 15 minutes at 37 °C followed by 15 minutes at 80 °C and the samples were purified using 1x AMPure XP beads (Beckman Coulter). For the Panhandle protocol, End-repair, dA tailing, native barcode ligation and sequencing library preparation were performed according to the Oxford Nanopore Technologies’ SQK-DCS109 kit (Direct cDNA Native Barcoding with EXP-NBD104 and EXP-NBD114). The fours samples were pooled at equal concentration and sequenced on a pre-used washed PromethION flow cell which had 4593. For both ONT and Panhandle protocols, we used 150 ng of prepared library to load on the flow cell. Sample concentration and profiles were determined using Qubit dsDNA HS 1X solution (Thermo Fischer scientific) and D5000 Tapestation (Agilent), respectively.

**Table 1:**
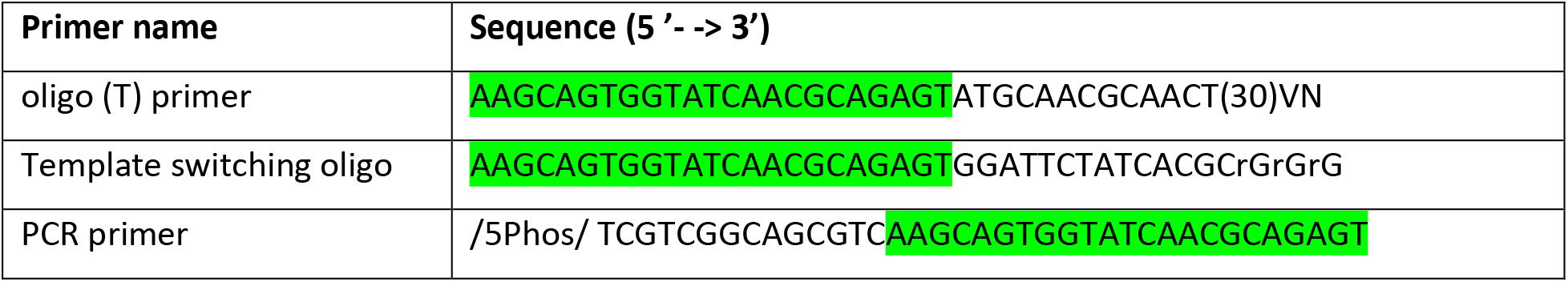
Primers used in this study. The common sequence across all primers is shaded in green.

### Basecalling, demultiplexing, and read processing

Reads were basecalled and demultiplexed during sequencing using Guppy (version 5.1.13) included within the MinKNOW suite package (version 21.11.7). We processed reads generated with ONT protocol using Pychopper (Oxford Nanopore Technologies) to both put them in the correct orientation and remove barcodes and sequencing adapters. Reads were then processed using Cutadapt (Martin, 2011) to remove poly(A) tails using the “-a “A[100]”” option. For reads generated using the Panhandle protocol, we used Pychopper to orient them and remove barcodes and sequencing adapters. We then used Porechop (Wick) to remove our custom added ITR primers and lastly used Cutadapt to remove poly(A) tails.

### Read alignment and coverage analysis

Medfly reads were aligned to the Medfly genome (Ccap_2.1, Genbank ID GCA_000347755.4, Refseq GCF_000347755.3). We used the NCBI *Ceratitis capitata* Annotation Release 103 for the transcriptome/genes/transcripts. Reads from cell lines MCF7, HeLa, and GM24143, which are all human-derived, were aligned to the recently published telomere-to-telomere T2T-CHM13 genome assembly, the first human gapless human genome completed with the help of long-read technologies (Nurk et al., 2022). We used the catLiftOffgenesV1 gene models which are GENCODE v35 gene models plus extra paralogs. These were generated using the Comparative Annotation Toolkit (CAT) (Fiddes et al., 2018) to lift over GENCODE models to T2T-CHM13 assembly and then using Liftoff (Shumate & Salzberg, 2020) to map genes missed by CAT and add other paralogs. CAT was also used to add gene models generated using PacBio Iso-Seq data. We selected the T2T-CHM13 assembly and the associated gene models as we believe they represent a more complete annotation of the human genome and transcriptome, respectively. All alignments were done using Minimap2 (Li, 2018) in splice-aware mode. In some cases, we subsampled reads using Seqtk (Li) before alignment. Alignment statistics were determined from bam files using Samtools (Li et al., 2009). To assess gene body coverage, we used RSeQC (Wang et al., 2012).

### Transcriptome construction and quality assessment

Flair (Tang et al., 2020) was used to construct a genome-guided transcriptome assembly from the three human cell lines, GM24143, MCF7, and HeLa. Briefly, 2 million PASS reads were subsampled using Seqtk for HeLa and GM24143 cell lines while for MCF7, we used all PASS reads available (3.4 million for ONT protocol and 2.5 million for Panhandle protocol). The six constructed transcriptomes were assessed using SQANTI (Tardaguila et al., 2018).

## Results

### Panhandle protocol shows higher cDNA-PCR amplicon yield

Following 0.8x AMPure XP magnetic beads cleanup of the PCR-amplified cDNA we used 1x dsDNA High Sensitivity Qubit kit (Thermo Fischer Scientific) to quantify the amount of amplicons yielded. We observed a higher yield of amplicons from the Panhandle method compared to the ONT method. From the 4 samples tested we obtained 1590, 1290, 1410, and 2086 ng of amplicons using the Panhandle method from MCF7, HeLa, GM24143, and Medfly, respectively, compared to 300, 576, 564, and 252, respectively using ONT method (**Table 2**, **Figure 2**). This is a range of 2.2 to 8.3-fold increase in yield (**Figure 2**, **Supplementary Figure 1**). Further, the cDNA profile of samples from Panhandle protocol showed a significantly reduced amount of molecules below 600 bp compared to ONT protocol.

**Table 2:** Total yield of PCR amplified cDNA. Total RNA from four samples was processed either following Oxford Nanopore Technologies’ SQK-PCB109 protocol or an in-house optimised protocol referred to as Panhandle. Following 20 cycles of PCR amplification, the yield of amplicons was measured using 1x dsDNA HS Quibit kit.

|  | ONT | Panhandle | Fold increase |
| --- | --- | --- | --- |
| MCF7 | 300 | 1590 | 5.3 |
| HeLa | 576 | 1290 | 2.2 |
| GM24143 | 564 | 1410 | 2.5 |
| Medfly | 252 | 2086 | 8.3 |

**Figure 2:**
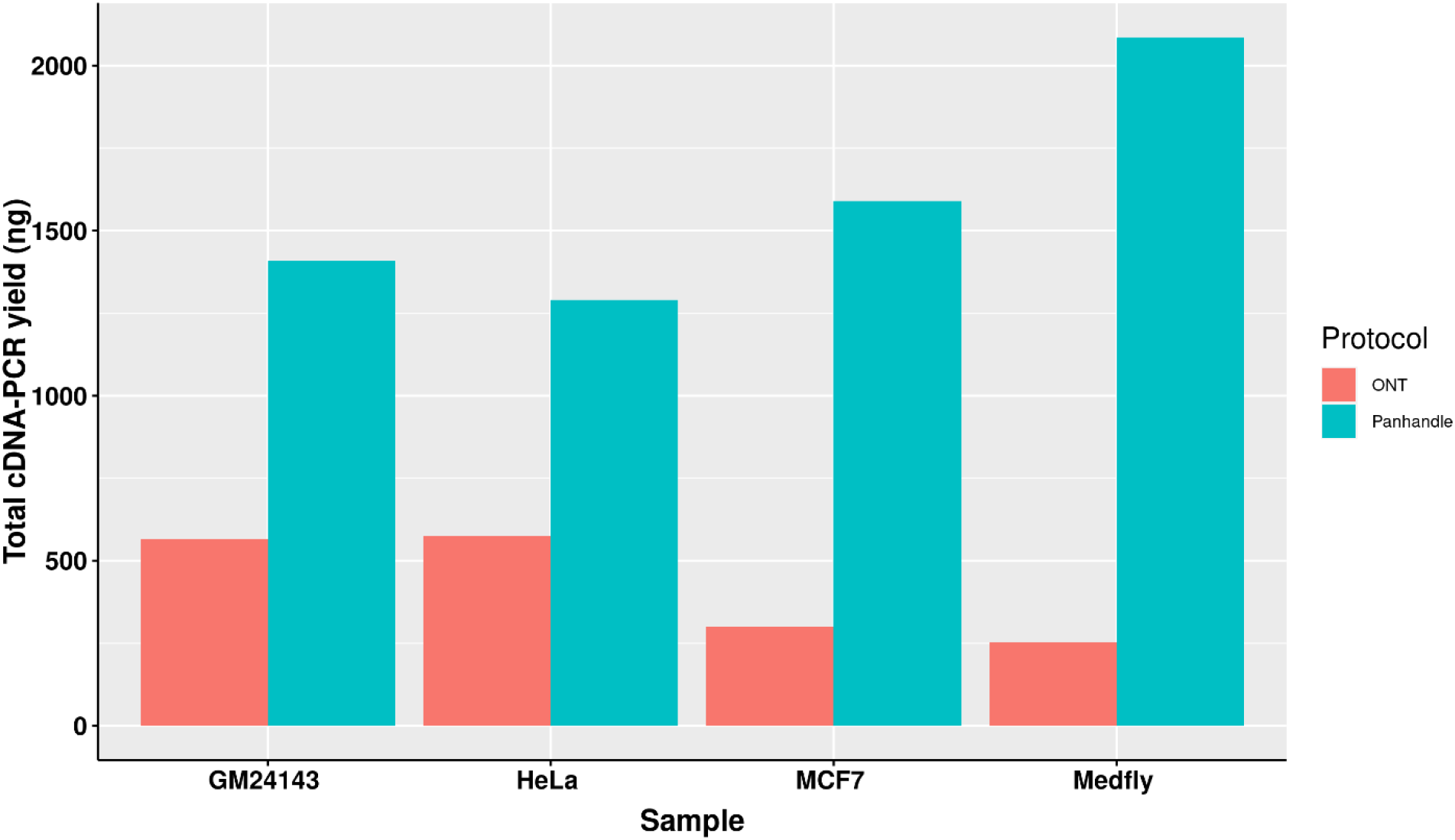
Total yield of PCR amplified cDNA. Total RNA from three human-derived cell lines and one from Mediterranean fruit fly embryos (Medfly) was reverse transcribed and amplified for 20 cycles either following the Oxford Nanopore Technologies’ SQK-PCB109 (ONT) or our in-house protocol called Panhandle and the yield of purified amplicons measured.

### Panhandle protocol yields longer reads

Samples prepared either with the ONT protocol or Panhandle protocol were pooled separately and sequenced on the PromethION using 2 separate flow cells, respectively. We obtained similar number of reads (**Supplementary Figure 2**). **Table 3** summarises the reads statistics. The ONT protocol samples yielded between 3.4 and 7.5 million reads while Panhandle protocol samples had between 3.2 and 9.8 million reads. Although 82 % and 72 % of the reads generated with the ONT protocol and Panhandle protocol, respectively were assigned to their respective barcode, we noticed a higher percentage of unclassified reads with the Panhandle protocol (28 % with Panhandle protocol versus 18 % with ONT protocol). Among higher quality reads referred to as PASS reads, the Panhandle protocol showed a much higher number of unclassified reads (9.8% with Panhandle protocol versus 1.4 % with ONT protocol, **Supplementary Figure 3**). The number of PASS and FAIL reads also seemed to differ. The Panhandle protocol seemed to yield higher number of PASS reads than the ONT protocol (**Supplementary Figure 4**), although we did not see this in another experiment.

**Table 3:** Sequenced read statistics. Four samples, processed either with ONT or Panhandle protocol, were sequenced on the PromethION. The number of Pass reads (reads with Phred score of 9 and above), Fail reads (reads with Phred score of 8 and below), total reads is shown.

| Sample | Protocol | Pass reads | Fail reads | Total sequenced reads |
| --- | --- | --- | --- | --- |
| MCF7 | ONT | 3,399,426 | 1,449,111 | 4,848,537 |
| HeLa | ONT | 2,487,353 | 974,869 | 3,462,222 |
| GM24143 | ONT | 3,432,169 | 1,455,416 | 4,887,585 |
| Medfly | ONT | 5,125,182 | 2,389,878 | 7,515,060 |
| Unclassified | ONT | 207,012 | 4,303,292 | 4,510,304 |
| Total_ONT | ONT | 14,651,142 | 10,572,566 | 25,223,708 |
| MCF7 | Panhandle | 2,601,122 | 670,034 | 3,271,156 |
| HeLa | Panhandle | 4,489,611 | 1,233,938 | 5,723,549 |
| GM24143 | Panhandle | 2,662,754 | 713,793 | 3,376,547 |
| Medfly | Panhandle | 7,923,177 | 1,927,617 | 9,850,794 |
| Unclassified | Panhandle | 1,919,040 | 6,830,558 | 8,749,598 |
| Total_Panh | Panhandle | 19,595,704 | 11,375,940 | 30,971,644 |

We compared read lengths after trimming Oxford Nanopore sequencing adapters, barcodes, our ITR adapters, and poly(A) tails. We consistently observed longer read lengths in the Panhandle protocol-generated reads compared to ONT protocol-generated reads (**Figure 3**, **Supplementary Figure 5**). Among the 3 human-derived cell lines, the 1^st^ quartile, median, mean, and 3^rd^ quartile where all either more than doubled or about doubled in the Panhandle protocol compared to the ONT protocol (207 vs 37, 454 vs 139, 537 vs 248, and 701 vs 379, respectively, **Table 4**).

**Figure 3:**
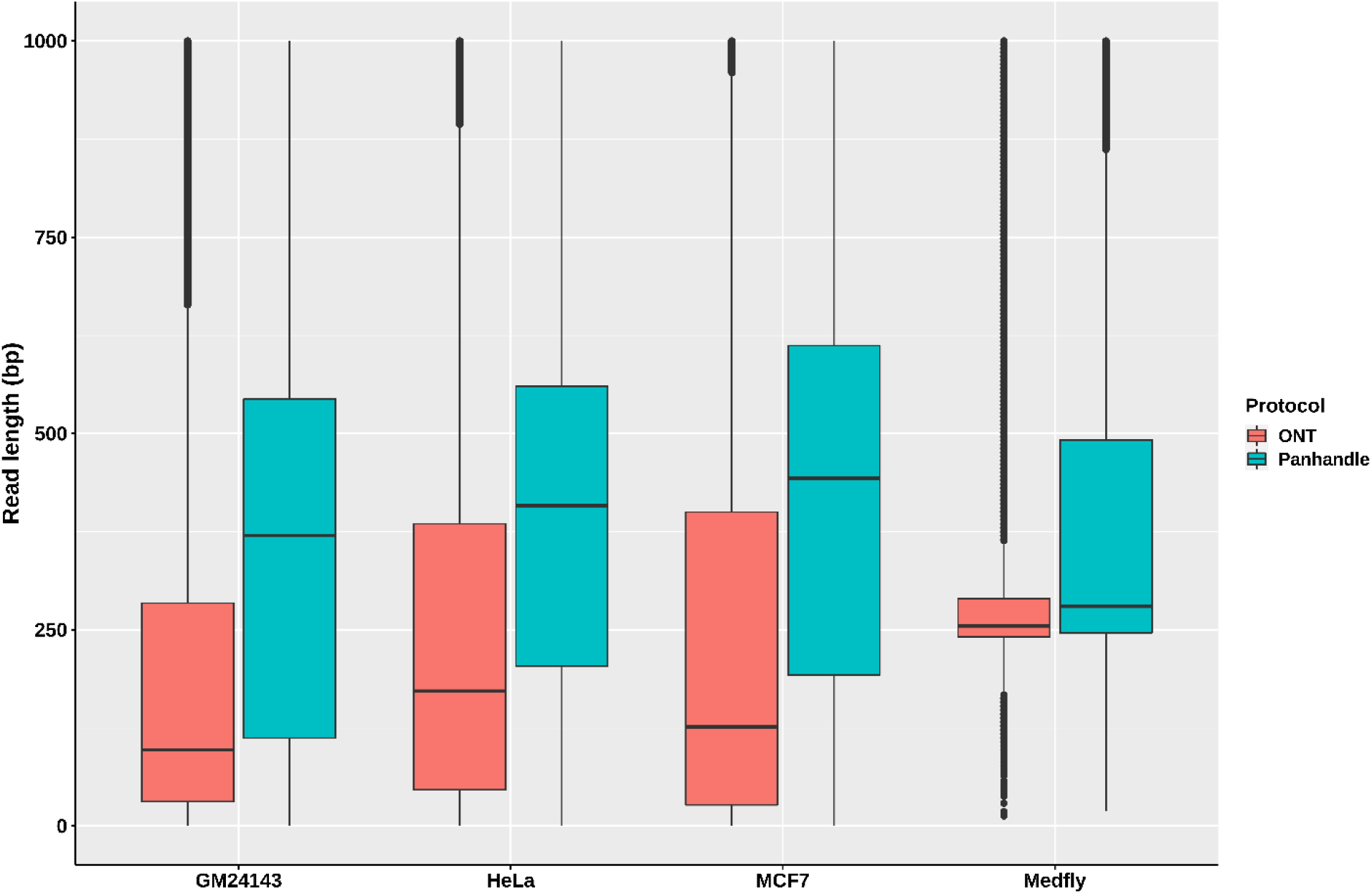
Read length distributions. Pass reads (reads with Phred score of 9 and above) from 4 samples processed both with ONT protocol (SQK-PCB109, Oxford Nanopore Technologies) and out in-house protocol called Panhandle were trimmed of all adapters and poly(A) tails. The boxplot shows their read length distributions.

**Table 4:** Processed Pass read statistics. Following cDNA-PCR sequencing on the PromethION of 4 samples processed either with ONT or Panhandle protocols, Pass reads were trimmed of all adapters and poly(A) tail and the distribution of their lengths measured. All presented figures are in basepairs.

| Sample | Protocol | 1<sup>st</sup> Quartile | Median | Mean | 3<sup>rd</sup> Quartile |
| --- | --- | --- | --- | --- | --- |
| GM24143 | ONT | 31 | 100 | 204.1 | 296 |
| HeLa | ONT | 48 | 178 | 256.6 | 399 |
| MCF7 | ONT | 31 | 139 | 284.3 | 441 |
| Medfly | ONT | 70 | 83 | 143.6 | 118 |
| GM24143 | Panhandle | 128 | 424 | 506.9 | 660 |
| HeLa | Panhandle | 224 | 437 | 491.4 | 633 |
| MCF7 | Panhandle | 269 | 500 | 613.5 | 810 |
| Medfly | Panhandle | 78 | 103 | 282 | 223 |

### Panhandle protocol shows higher genome alignment rate

We used Seqtk to subsample one million reads from each sample and aligned it to the respective transcriptome using Minimap2. We then assessed alignment rates using Samtools. We observed 19 – 28 % higher alignment rate with Panhandle protocol-generated data compared to ONT protocol-generated data.

### Panhandle protocol shows better gene body coverage

We aligned reads generated from each sample to their respective genomes and assessed gene body coverage using RSeQC (Wang et al., 2012). We noticed a marked increase in gene body coverage (uniformity of coverage) with reads generated with the Panhandle protocol. Reads generated with the ONT protocol showed a marked 3′ bias with only about 40 – 50 % of reads showing full-length coverage of the genes (**Figure 4**). Reads generated using the Panhandle protocol showed slight 5′ bias except for Medfly reads which had a more marked 5′ bias.

**Figure 4:**
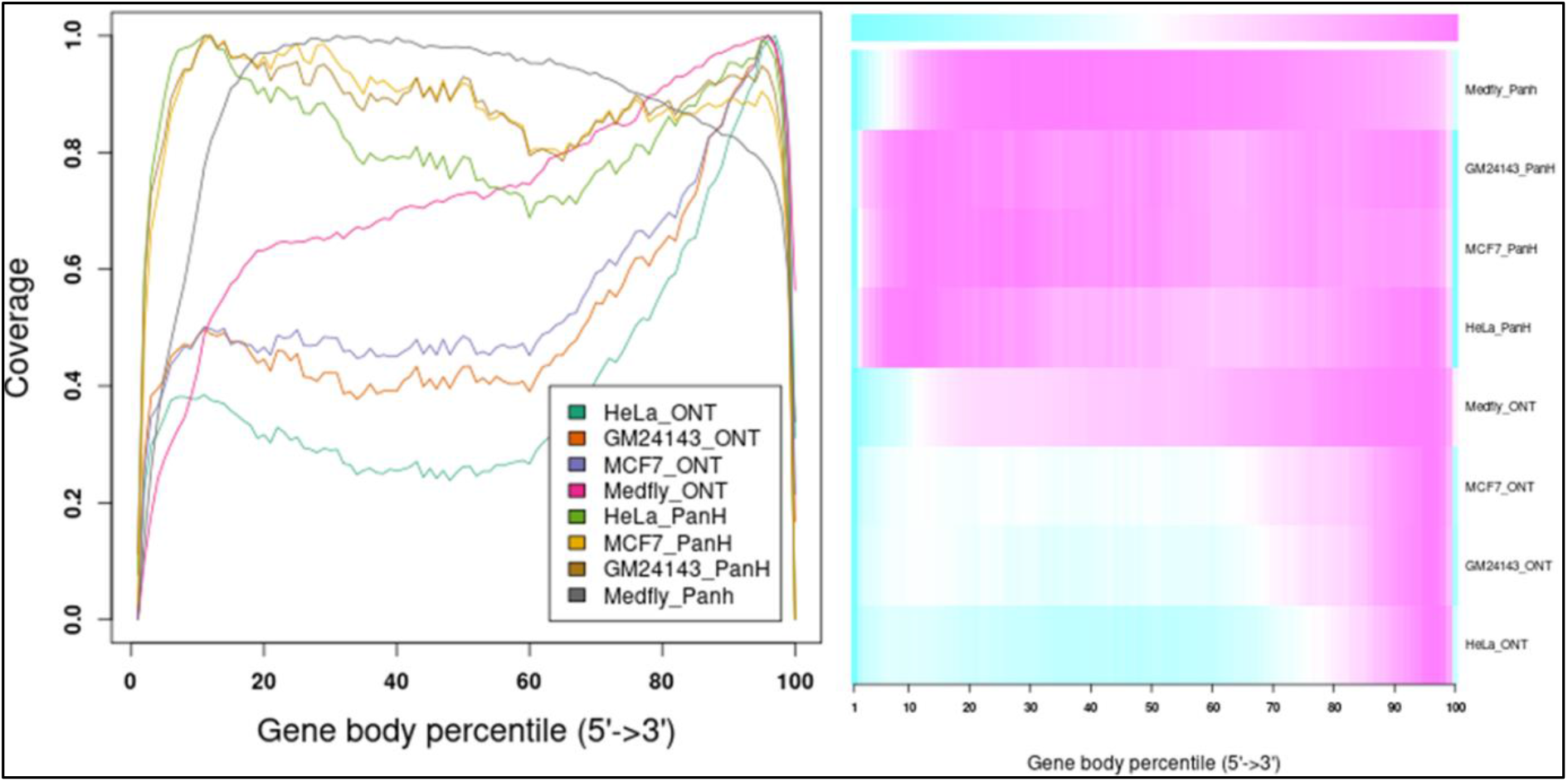
Gene body coverage comparison between ONT and Panhandle protocols. Total RNA from three human-derived cell lines (MCF7, HeLa, and GM24143, and Mediterranean fruit fly embryos) was processed both with the ONT protocol (SQK-PCB109, Oxford Nanopore Technologies) and our in-house optimised protocol called Panhandle. One million subsampled Pass adapter and poly(A) tailed trimmed reads were aligned to the respective genomes and gene body coverage assessed using RSeQC (Wang et al., 2012). A) Line graph showing gene body coverage for all four samples processed either with Oxford Nanopore Technologies’ (ONT) SQK-PCB109 protocol or our in-house Panhandle protocol (PanH or Panh). B) Same data used in ‘A’ but represented as a Heat map.

We further obtained data previously generated using the MCF7 cell line (Chen et al., 2021). The authors followed manufacturer’s instructions and performed direct cDNA sequencing (without PCR amplification, here-in referred to as cDNA-direct), direct RNA sequencing (RNA-direct), and sequencing of PCR amplified cDNA (cDNA-PCR). We compared gene body coverage obtained by the authors to our data generated using Panhandle protocol (**Figure 5**). We observed 3′ bias in the authors’ data particularly with cDNA-PCR dataset. Overall, our data showed better gene body coverage particularly at the 5′ whereas the authors’ data showed better coverage at the extreme 3′ end. Still, over 80 % of the reads generated with the Panhandle protocol show near full-length gene coverage.

**Figure 5:**
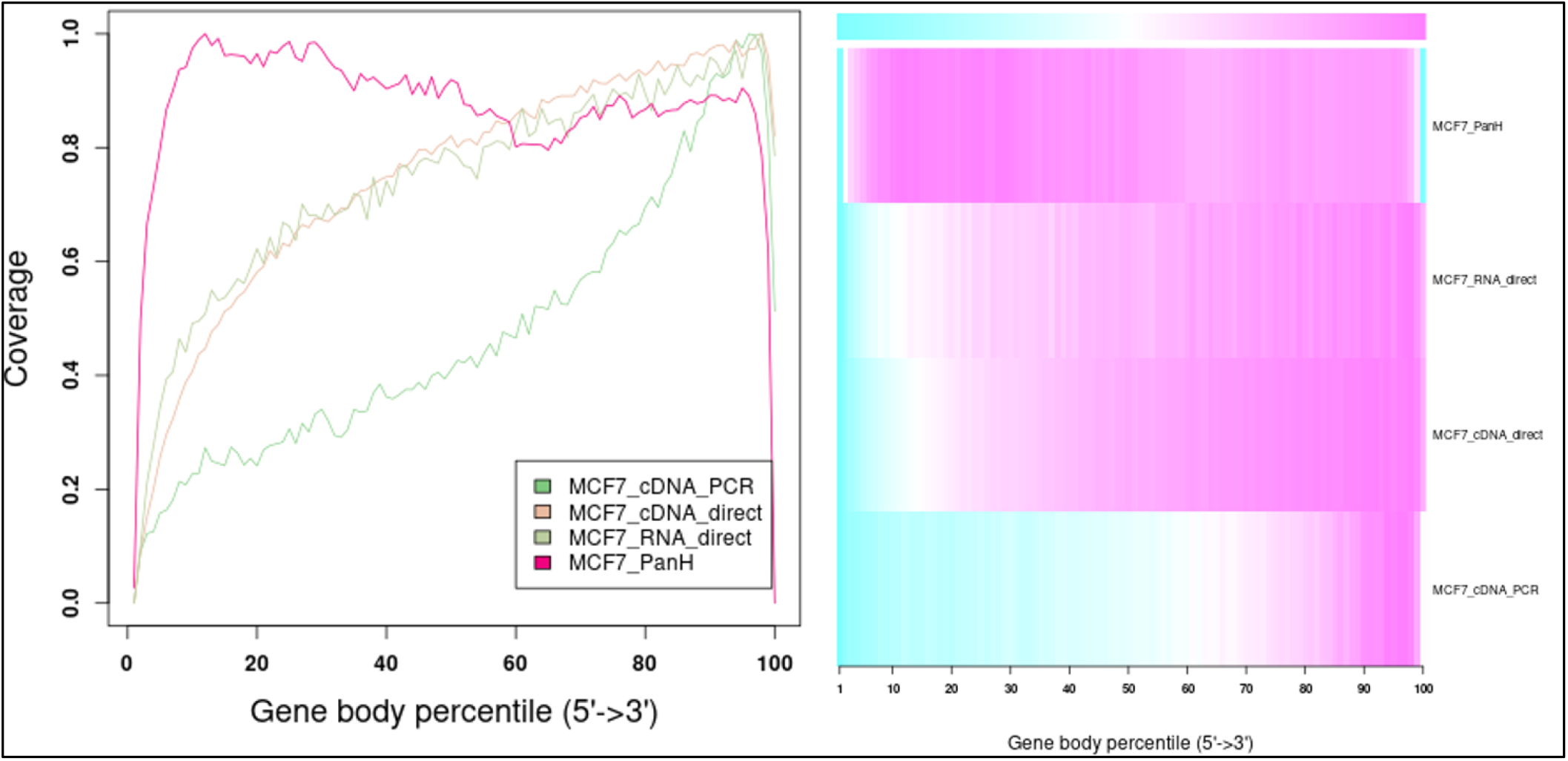
MCF7 Gene body coverage comparison between Panhandle protocol and previously published datasets. We downloaded previously published MCF7 datasets (Chen et al., 2021) that were sequenced through cDNA-PCR, direct cDNA sequencing (cDNA-direct), and direct RNA sequencing (RNA-direct) following Oxford Nanopore Techologies protocols. We aligned these reads to the T2T-CHM genome together with our MCF7 dataset generated through our Panhandle protocol and assessed gene body coverage using RSeQC (Wang et al., 2012). A) Line graph showing gene body coverage using MCF7 cell line and comparing our inhouse protocol called Panhandle (PanH) with other Oxford Nanopore Technologies (ONT) RNA sequencing approaches. B) Same data used in ‘A’ but represented as a Heat map.

### Panhandle protocol yields higher quality long-read transcriptome assembly

We constructed the transcriptomes of three cell lines; MCF7, HeLa, and GM24143 using Flair and then assessed the transcriptomes using SQANTI. The Panhandle protocol consistently generated a higher number of genes that matched annotated genes. For example, the Panhandle protocol generated 10226, 10525, and 11660 genes matching annotated genes compared to 7780, 9955, and 11126 genes from the ONT protocol, for GM24143, HeLa, and MCF7 cell lines, respectively. On the other hand, the ONT protocol generate over twice the number of novel genes than the Panhandle protocol (**Figure 6**A). Further, the Panhandle protocol generated almost double the percentage of transcripts with a full splice match (FSM) to annotated genes than the ONT protocol (average of 14.0 versus 7.2 %, respectively, **Figure 6**B). The number of genes with six or more isoforms was significantly higher in the Panhandle protocol compared to ONT protocol (Wilcox test p-value 0.08, **Figure 6**C). We also compared distance to annotated transcription start site (TSS) of constructed transcripts that showed incomplete splice match to their associated annotated transcript. We observed more restriction around the TSS in the Panhandle transcriptome than ONT transcriptome (**Figure 6**D).

**Figure 6:**
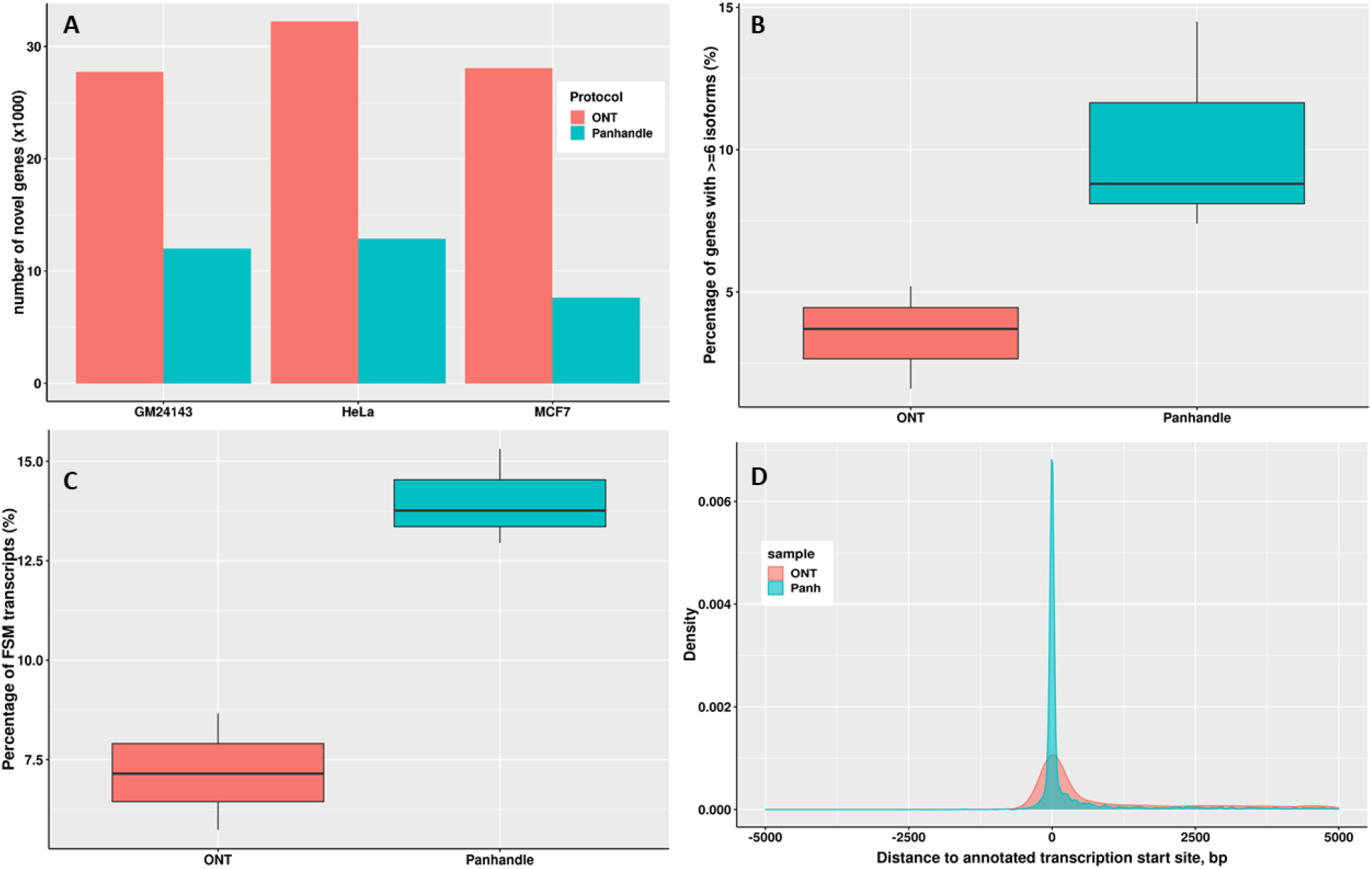
Transcriptome comparison between ONT and Panhandle protocols. We used Flair (Tang et al., 2020) to construct a genome-guided transcriptome of 3 cells line; GM24143, HeLa, and MCF7 using reads generated from either Oxford Nanopore Technologies (ONT) protocol SQK-PCB109 or our in-house protocol named Panhandle (or Panh, see Supplementary Protocol). The six transcriptomes constructed were assessed using SQANTI (Tardaguila et al., 2018). **A**) Total number of novel genes identified in each transcriptome. **B**) Percentage of genes with six or more isoforms. **C**) Percentage of transcripts whose splice-pattern (exon structure) is a complete match or full-splice match (FSM) to the GENCODE v35 annotation liftoff to T2T-CHM13 genome assembly (Nurk et al., 2022). **D**) Distance of constructed transcripts’ transcriptional start site (TSS) to their associated annotated transcript TSS. We used MCF7 constructed transcripts in the incomplete splice match (ISM) category.

## Discussion and conclusion

Accurately sequencing full-length transcripts remains a main goal of RNAseq but still represents a challenge. As we show in Figure 4, long-read sequencing technologies such as Oxford Nanopore Technologies show 3′ bias for the majority of reads. A method has been previously suggested to improve enrichment of longer molecules (Lukyanov et al., 1995). This method involves addition of an inverted terminal repeat in the cDNA synthesis primers such that a single primer is used during PCR. The ITR mediates the creation of panhandle-like structures during the annealing phase of each PCR cycle. These panhandle-like structures are more stable for short fragments compared to longer fragments and prevent primer binding in short molecules allowing longer molecules to be generated (Dai et al., 2007; Lukyanov et al., 1995). In the current study we adapted this method to Nanopore cDNA library preparation and implemented it on a variety of RNAs to show that it improves cDNA PCR amplicons yield by at least 2-fold and also greatly improves gene body coverage compared to the standard Nanopore cDNA library preparation protocol.

The Panhandle method resulted in a 2 – 8-fold increase in amount of PCR amplicons generated. This suggested improved PCR efficiency probably emanating from the suppression of primer binding in primer dimers and other short fragments. This is a very attractive attribute for samples with limited amount of total RNA, such as single cells and single embryos, as it increases sensitivity of PCR. Further, this attribute should improve overall RNAseq data quality as it potentially allows one to lower the number of PCR cycles needed to obtain ample amplicons for sequencing. Lowering the number of PCR cycles should result in less duplicates, polymerase errors, and reduced PCR bias towards shorter molecules.

Reads generated from the Panhandle protocol showed a higher alignment rate than ONT protocol. This is probably attributable to the longer reads. We observed that non-aligned reads were more enriched with shorter reads compared to aligned reads (**Supplementary Figure 6**).

Gene body coverage was our focus. The Panhandle protocol dramatically improved gene body coverage when compared to direct RNA sequencing, direct cDNA sequencing, and cDNA-PCR sequencing. It achieved a main goal of long-read RNAseq to obtain full-length transcripts for the majority of reads. Further optimisation of this method could improve gene body coverage. The Panhandle method showed reduced 3′ coverage compared to 5′ coverage and perhaps this can further be resolved in future studies. Piao et al. (Piao et al., 2001) discovered that ligation (or addition) of a long (35 bp) linker to a cDNA library followed by PCR amplification of the library using a short primer (17 bp) primer that is part of the linker yields long-transcript enriched libraries that are more representative of full-length cDNA libraries. This design suppressed the amplification of short fragments at the cost of long fragments. We have not tried this format but it might further improve gene body coverage. However, even as implemented in the current study, the Panhandle method reduces sequencing costs, improves yields which could increase sensitivity and reduce PCR artifacts, and yields better coverage across gene body. Although we did not specifically investigate the effect of the Panhandle method on gene expression quantification, improvement in data full-length data quality should lead to improved gene, and especially isoform, expression quantification.

The Panhandle protocol showed some drawbacks. It seemed to yield a smaller number of reads than the ONT protocol (**Supplementary Figure 7**). This is most likely attributable to sequencing adapter ligation method we used. We employed the enzymatic ligation method as detailed in the SQK-DCS109 protocol. The current SQK-PCB109 protocol employs a much more efficient ligation method based on click chemistry (Jaworski & Routh, 2018). In our hands, this method resulted in 2-fold increase in yields (**Supplementary Figure 8**). Further, we observed a higher percentage of unclassified reads among Panhandle protocol-generated reads. Among all reads, we observed a ~ 30 % increase in number of unclassified reads in the Panhandle protocol compared to standard ONT protocol. Among high quality PASS reads, we observed a 7-fold increase in unclassified reads in the Panhandle protocol compared to standard ONT protocol. Again, this is most likely attributable to the less efficient enzymatic ligation of barcodes. Barcodes in the ONT protocol are added to each sample through PCR while in the Panhandle protocol barcodes are enzymatically ligated. We have observed a higher number of unclassified reads in other samples were barcodes are enzymatically ligated following the SQK-LSK109 kit protocol. This is however, should be less worrisome as unclassified reads are enriched with shorter and lower quality reads. Further, the alignment rate of Panhandle protocol-generated reads is 19 - 28 % higher than ONT protocol which compensates for some of the reads lost due to being unclassified. For samples that are not barcoded no unclassified reads are expected. The remaining drawback of the ligation method we applied would be the lower number of sequenced reads. However, it should be possible for Oxford Nanopore Technologies to avail a ‘click chemistry’ version of the Panhandle protocol and thereby prevent loss of reads. Although we observed a higher rate of FAIL reads in the ONT protocol-generated reads we think this is related more to the quality of flow cell used as we did not see phenomenon is a separate experiment (**Supplementary Figure 9**).

Long-read RNAseq has been generally regarded as sequencing transcripts in their entirety. Therefore, unlike in short-read RNAseq where a transcriptome is assembled by identifying overlaps between reads, mostly commonly through de Bruijn graph-based algorithms (Grabherr et al., 2011), long-read transcriptomes are constructed by identifying reads emanating from the same genomic locus, either by alignment of reads to the genome (Kovaka et al., 2019; Tang et al., 2020) or self-alignment of reads (Li et al., 2017; Nip et al., 2020), and then collapsing all reads into a non-redundant set of genes and isoforms. However, as we show in **Figure 4** and **Figure 5** for cDNA-PCR method which is the most commonly used method of Nanopore RNAseq, over half of the reads generated do not cover the entire transcript. This, we think, stems from an intrinsic inefficiency in the PCR. Unfortunately, however, this potentially creates artifacts that can be potentially misconstrued as novel genes and/or alternative isoforms. Consequently, transcriptomes generated from all three cell lines using ONT protocol resulted in a higher number of novel genes but a lesser number of genes matching annotated genes and a lesser number of transcripts with a full-splice match to annotated transcripts. Further, we think that the Panhandle protocol generated more genes with six or more isoforms than the ONT protocol due to the longer reads generated with the Panhandle protocol. Although the Wilcox test p-value was high at 0.08, meaning the test are significant at 90% confidence interval, we think this is due to the sample size of three. Nevertheless, these attributes make the Panhandle protocol superior to the ONT protocol in generating higher quality transcriptomes.

An alternative approach is to enrich for full-length transcripts using cap dependent linker ligation (TeloPrime, Lexogen) combined with size selection (Cartolano et al., 2016), a method applied in conjunction with PacBio sequencing. Firstly, using 0.6X AMPure bead size selection did not improve ONT protocol (**Supplementary Figure 10**). Further, the Panhandle protocol described here is simpler and does not rely on a specific commercial kit. We strongly believe that the Panhandle protocol yields better results than the current ONT protocol. Although we used different cDNA synthesis and PCR amplification thermocycler conditions, we obtain similar results just by using the Panhandle primer design and following the ONT protocol. We used conditions described in the current study mostly as a precautionary measure since this is a lab-adapted protocol that has worked well over many samples.

## Supporting information

Supplementary protocol

## Acknowledgments

We would like to thank Professor Kostas Mathiopoulos, Laboratory of Genomics and Molecular Biology, Department of Biochemistry & Biotechnology, University of Thessaly, Thessaly, Greece, for providing the Mediterranean fruit fly embryo samples.

## Author contribution statement

AB conceived the idea, conducted the experiments, performed data analysis, and drafted the manuscript. SO performed data analysis, was involved in experimental design and consultation, and was involved in earlier Panhandle protocol development and optimisation. YCW was involved in earlier Panhandle protocol development and optimisation. KM was involved in experimental design. JR oversaw the project execution. All authors read and corrected the manuscript.

## Ethics statements

**Studies involving animal subjects:** No animals were involved in this study.

**Studies involving human subjects:** No human studies are presented in this manuscript.

**Inclusion of identifiable human data:** No potentially identifiable human images or data is presented in this study.

## Data availability statement

The datasets presented in this study have been deposited to NCBI’s SRA with project number PRJNA867697. The individual SRR numbers for each dataset can be found in Supplementary Table 1.

