## Supplementary Figures for "Improved Nanopore full-length cDNA sequencing by PCR-suppression"

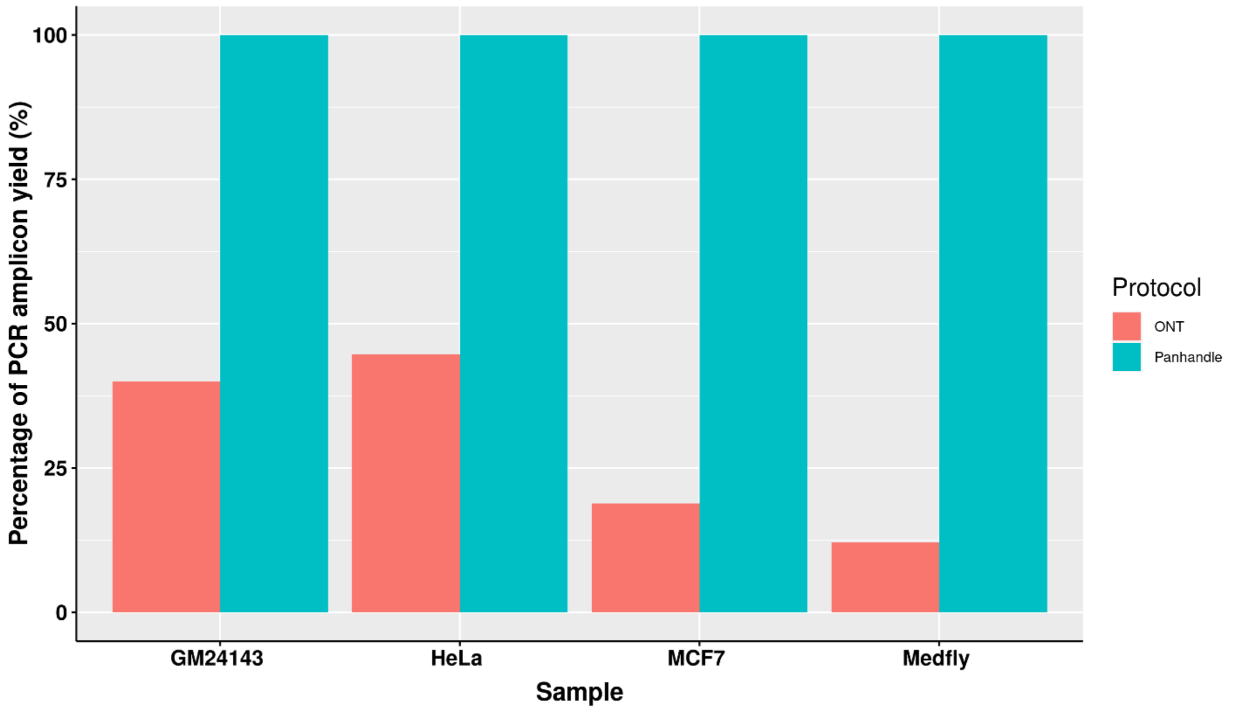


**Supplementary Figure 1**: Same as **Figure 2** but showing the yield from ONT protocol as a percentage of the yield from Panhandle protocol.

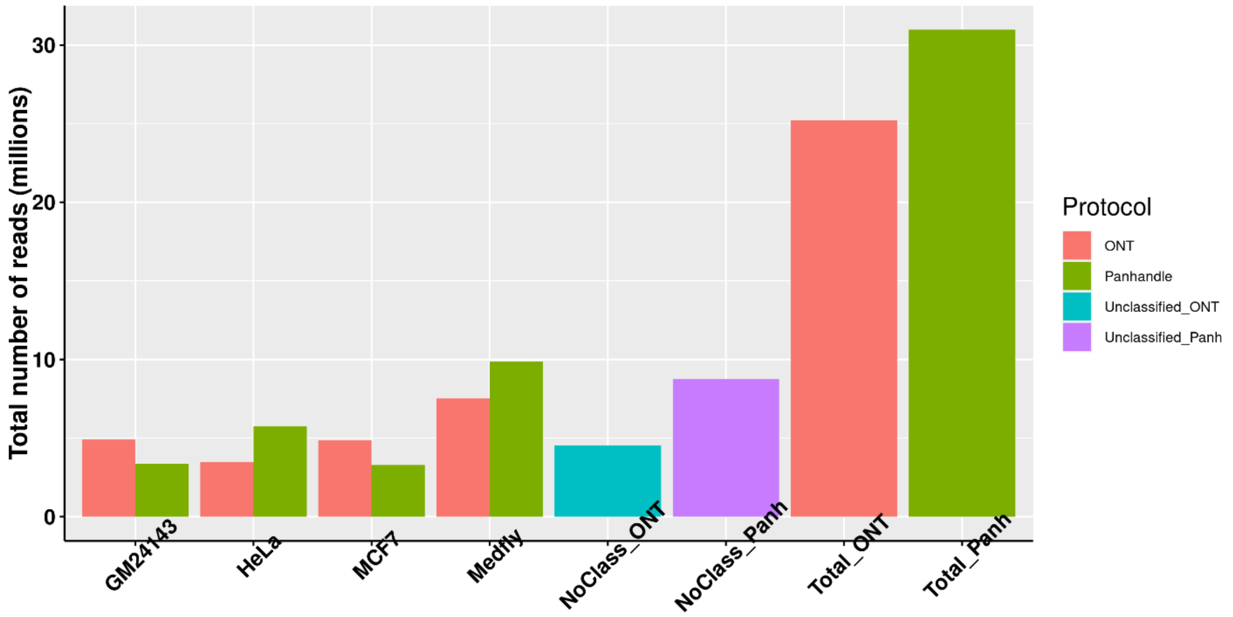


**Supplementary Figure 2: Total sequenced read numbers**. The total number of reads generated from each of the 3 human-derived cell lines (GM24143, HeLa, and MCF7) and Mediterranean fruit fly embryos (Medfly) is shown. Samples were processed either with Oxford Nanopore Technologies' SQK-PCB109 (ONT) or our in-house protocol (Panhandle). Samples processed with the sample protocol were barcoded, pooled and sequenced on the same PromethION flow cell. Following sequencing, reads were demultiplexed. The number of reads that were not successfully assigned to a correct barcode are shown with a prefix 'NoClass' for each protocol. The total number of sequenced reads is also shown for each protocol.

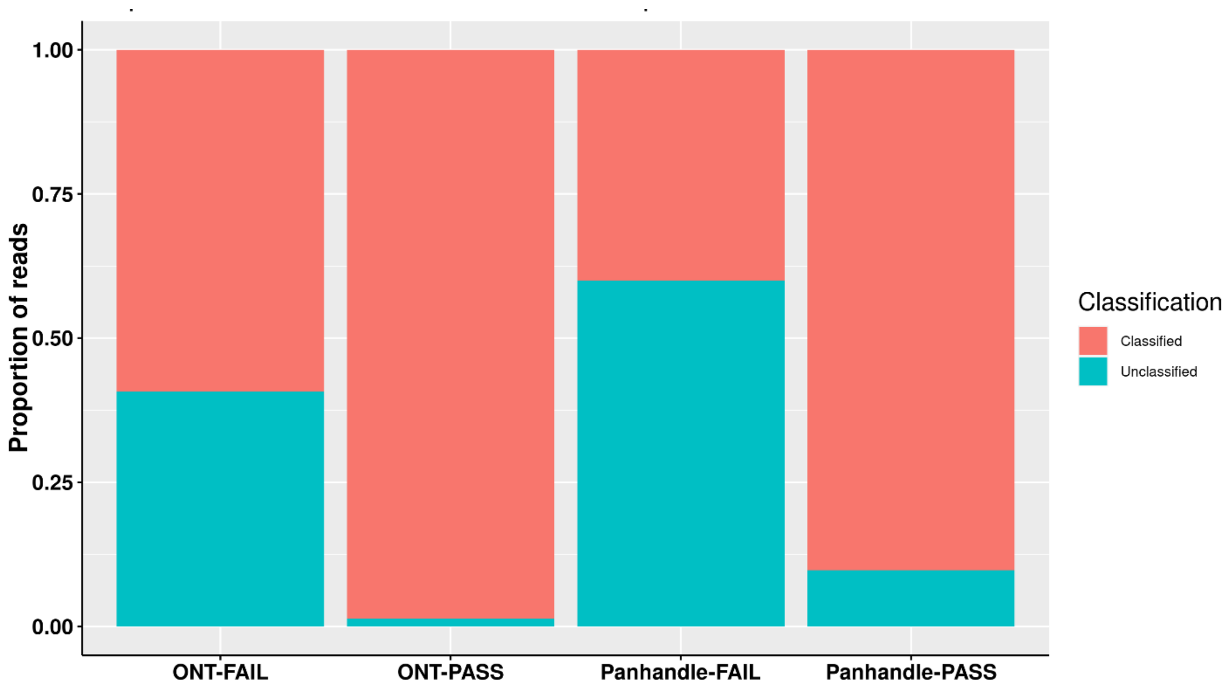


**Supplementary Figure 3: Proportion of classified reads**. Four samples were processed both with Oxford Nanopore Technologies' SQK-PCB109 protocol (ONT) and our in-house protocol called Panhandle. ONT and Panhandle processed samples were barcoded, pooled, and sequenced on different PromethION flow cells. The number of reads successfully assigned to their barcode of origin (Classified) and those unassigned to their correct barcode (Unclassified) from both Pass reads (reads with Phred score of 9 and above) and Fail reads (reads with Phred score of 8 and below) are shown.

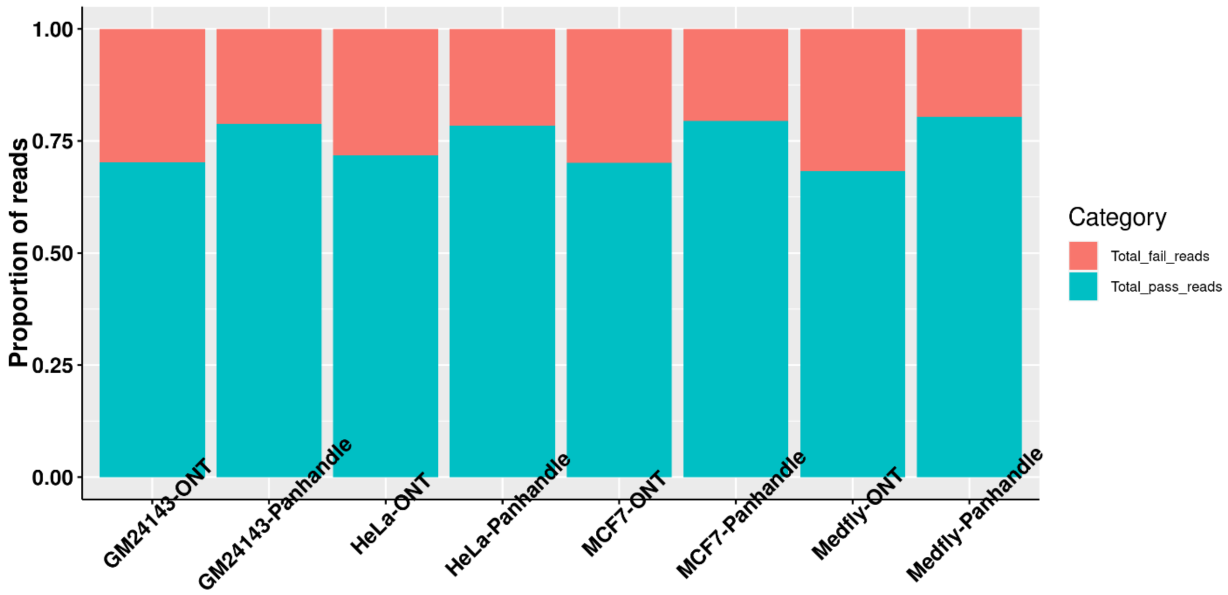


**Supplementary Figure 4: Proportion of Pass and Fail reads**. Same as **Figure 5** but showing proportion of Pass and Fail reads for each sample as a total of the number of reads for that sample.

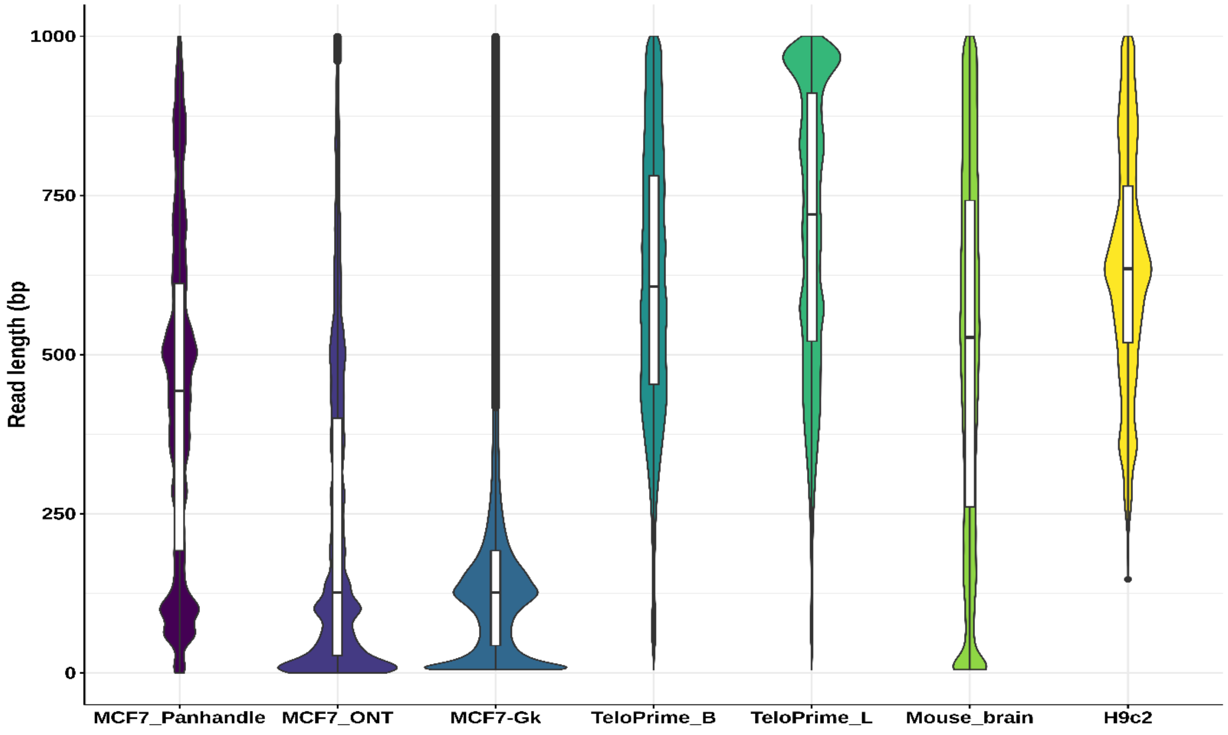


**Supplementary Figure 5: Read length distributions**. The length of sequenced reads from seven different experiments are compared. MCF_Panhandle and MCF7_ONT are reads generated in the current work from MCF7 cell line using either our in-house protocol called Panhandle or Oxford Nanopore Technologies' (ONT) SQK-PCB109 protocol, respectively. MCF7-Gk are reads generated by Göke lab from MCF7 cell line following ONT's SQK-PCS108 (Chen et al., 2021). TeloPrime_B and TeloPrime_L are reads generated from mouse brain and liver tissues, respectively and processed following TeloPrime Full-Length cDNA

Amplification protocol (Lexogen) followed by ONT's SQK-LSK108 protocol (Sessegolo et al., 2019). Mouse_brain are reads generated from mouse brain tissue and processed following ONT's SQK-LSK108 protocol (Sessegolo et al., 2019). H9c2 are reads generated from embryonic heart derived H9c2 cell line and processed following ONT's SQK-PCS108 protocol (Cao et al., 2021). All samples here were derived from PCR-amplified cDNA.

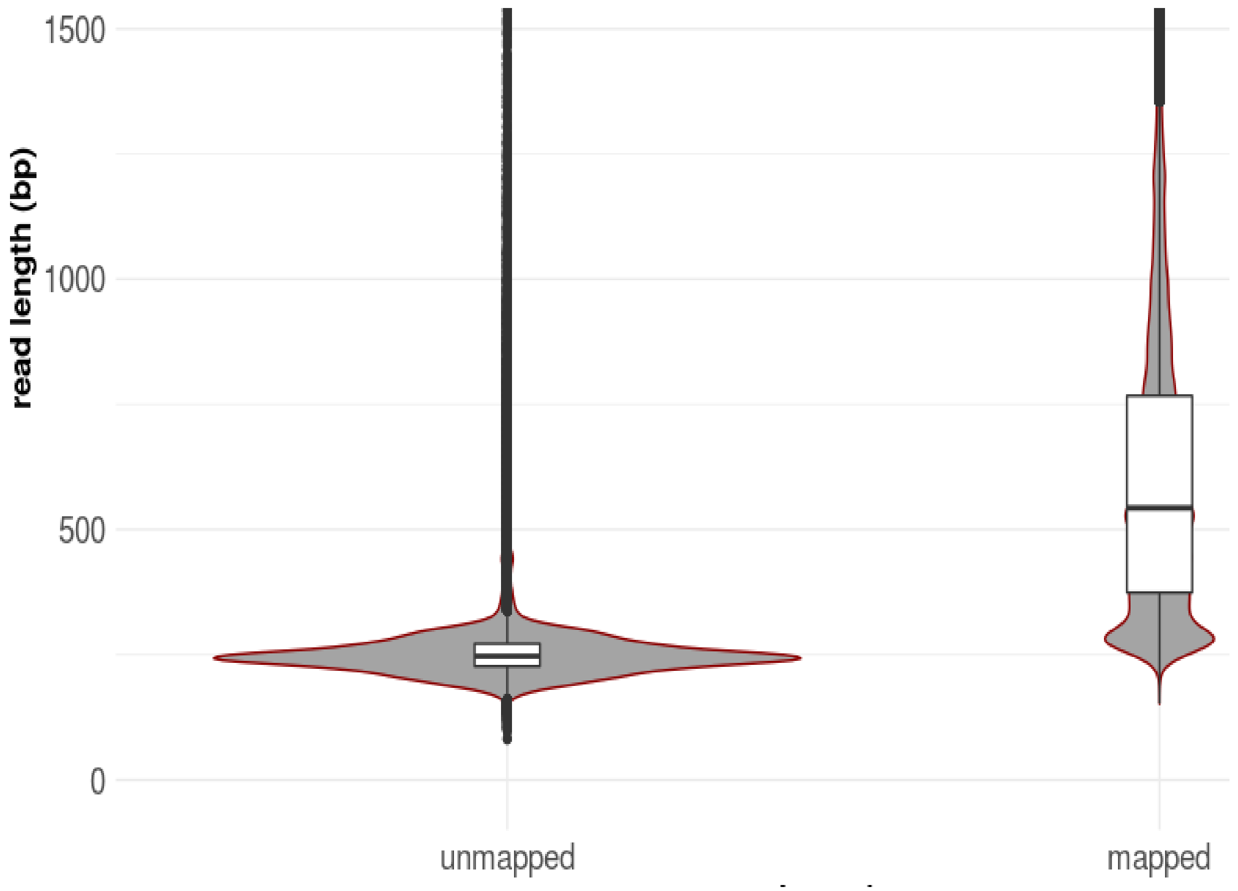


**Supplementary Figure 6: Read length of mapped and unmapped reads**. Reads from Medfly sample were aligned to the genome (Ccap_2.1, Genbank ID GCA_000347755.4, Refseq GCF_000347755.3). The length of reads mapped and unmapped to the genome were compared.

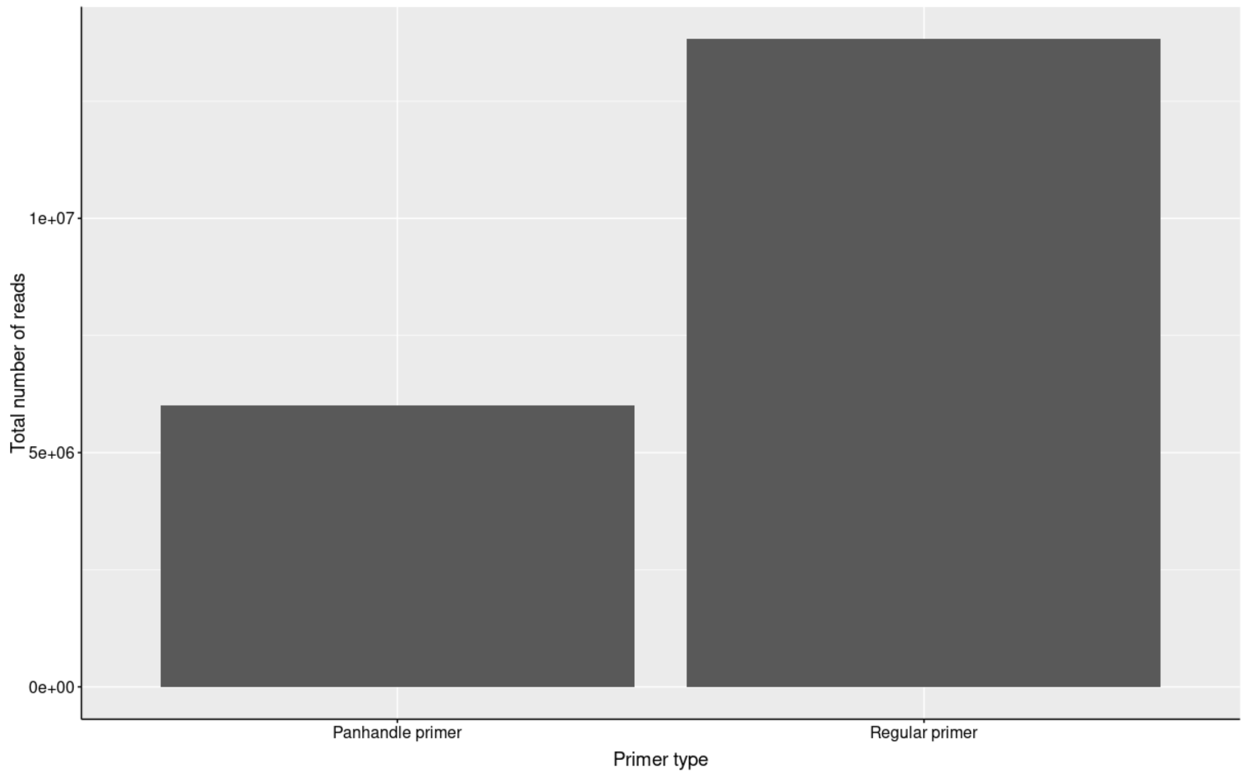


**Supplementary Figure 7: Total number of reads generated per protocol.** Number of pores at QC were 1695 and 737 for Panhandle and ONT primers, respectively. Samples were sequenced on 2 separate previously used PromethION flow cells.

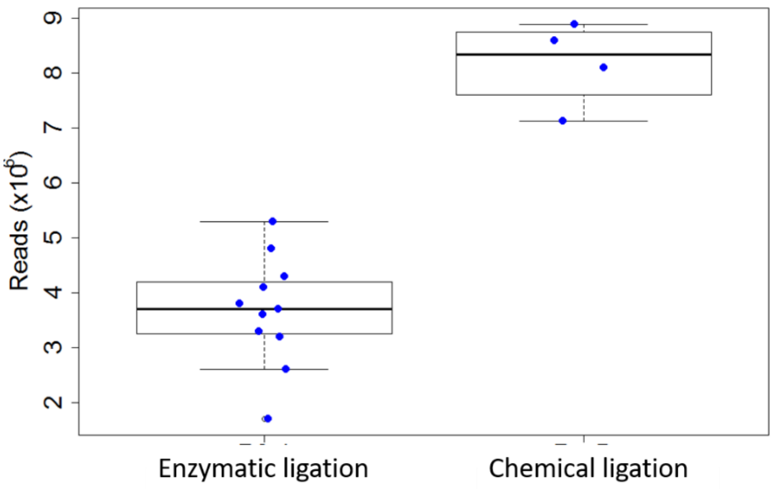


**Supplementary Figure 8: Sequencing throughput comparison of Enzymatic and chemical sequencing adaptor ligation.** Total RNA from Mediterranean fruit fly embryos was processed either with our in-house Panhandle protocol or with Nanopore SQK-PCS108 protocols and the libraries sequenced on MinION. The number of generated reads is shown here. Our in-house Panhandle protocol employs enzymatic ligation of sequencing adapters while the SQK-PCS108 protocol employs 'click chemistry' mediated ligation of sequencing adaptors.

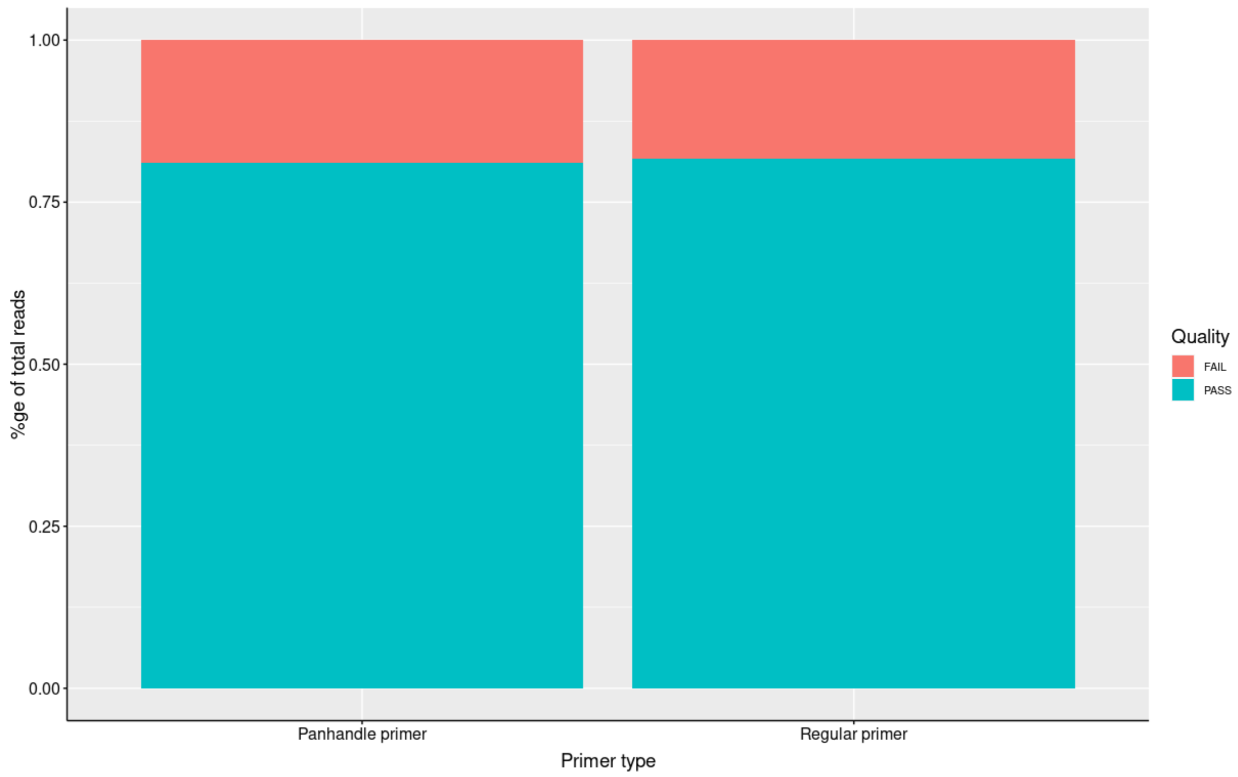


**Supplementary Figure 9: Percentage of PASS and FAIL reads per protocol.** Primers/protocol don't seem to influence barcode assignment

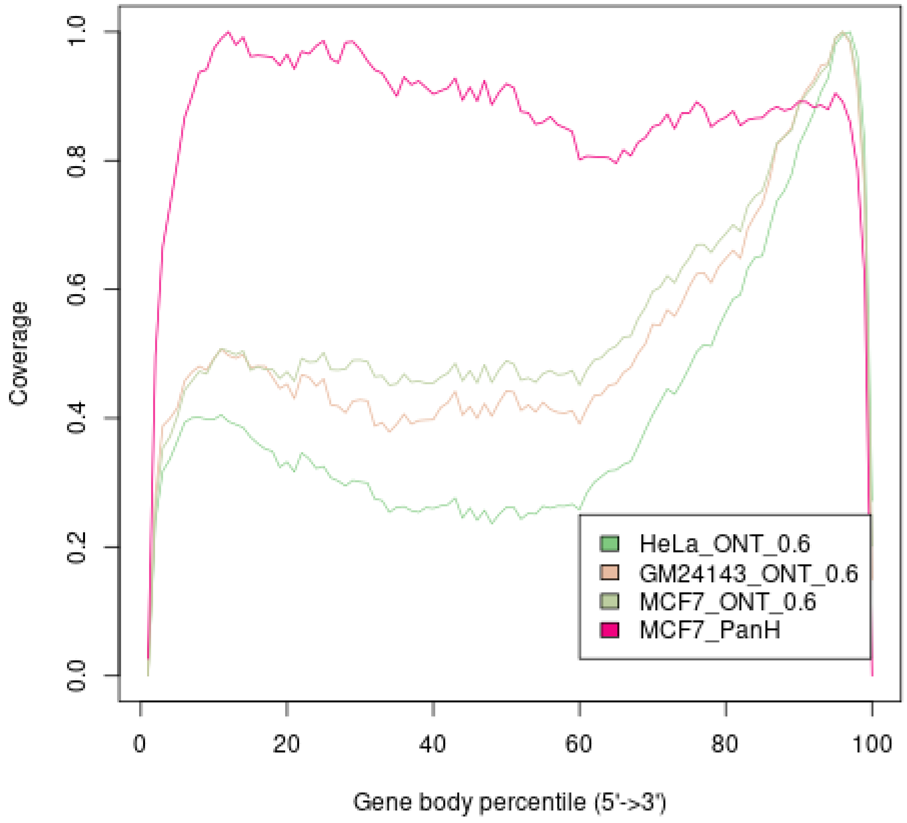


**Supplementary Figure 10**: **Gene body coverage comparison between Panhandle protocol and ONT protocol with 0.6X AMPure size-selection**. Total RNA from three human-derived cell lines (MCF7, HeLa, and GM24143) was processed both with the ONT protocol (SQK-PCB109, Oxford Nanopore Technologies) and our in-house optimised protocol called Panhandle. For the ONT protocol, we performed a 0.6X AMPure XP beads size-selection on the pooled library. Libraries were sequenced on the PromethION. One million subsampled Pass adapter and poly(A) tailed trimmed reads were aligned to the respective genomes and gene body coverage assessed using RSeQC (Wang et al., 2012).
