## Supplementary protocol for "Improved Nanopore full-length cDNA sequencing by PCR-suppression"

**Purpose:**

1. Compare Panhandle primers to Nanopore primers
2. Thaw and make the following.
   a. BioChain MCF, make up to 100 ng/µl
   b. HeLa total RNA (10 ng/µl)
   c. GM24143 total RNA (83 ng/µl)
   d. Cc-E-6H total RNA (100 ng/µl).
3. Perform 2 protocols;
   a. Our in-house protocol
   b. ONT PCB109.

Perform Nanopore Barcoded cDNA amplification and library preparation.
Compare cDNA profiles and yields

**Process Overview:** Poly-A tailed mRNA is reverse transcribed to cDNA. The synthesized cDNA is then amplified, followed by validation of its electropherogram using Caliper LabChip instrument or Tapestation, and its yield is quantified with a Qubit HS DNA assay.

**Materials**

..

**170522**
**B002-05-6C**

**Nanopore protocol: B002-05-6C1**

1. Thaw the RT and PCR reagents.
2. Setup the Nanopore protocol RNA-Primer annealing reaction as follows:

| | | A | B | C | D | -Ve |
|---|---|---|---|---|---|---|
| | | **MCF7** | **HeLa** | **GM24143** | **Cc-E-6H** | **Water** |
| No. | **Component** | **volume/sample (µl)** | **vol (µl)** | **vol (µl)** | **vol (µl)** | **vol (µl)** |
| 1. | dNTP | 1 | 1 | 1 | 1 | 1 |
| 2. | Water | 8 | 4 | 8 | 8.5 | 9 |
| 3. | Oligo-dT (VN primer) | 1 | 1 | 1 | 1 | 1 |
| 4. | Template RNA | 1 | 5 | 1 | 0.5 | 1 |
| | **Total Volume** | **11** | **11** | **11** | **11** | **11** |

I will follow PCB109 protocol as much as possible.

3. Mix gently and spin down briefly.
4. Heat at 65 °C for 5:00 mins, and quickly chill in ice/water bath for at least 1:00 min
5. Setup Reverse Transcription (RT) mastermix as follows:

| **RT mix** | | **x samples** |
|---|---|---|
| **Component** | **Volume/sample (µl)** | **5** |
| Water | 1 | 5 |
| Maxima H 5X Buffer | 4 | 20 |
| RNase inhibitor | 1 | 5 |
| TSO (10 uM) | 2 | 10 |
| **Total Volume** | **8** | |

6. Mix gently by flicking the tube then spin down.
7. Add the 8 µl of the RT mix to the 11 µl annealed template RNA sample.
8. Incubate at 42 °C for 2 minutes
9. Add 1 ul of Maxima H RTase for a total reaction volume of 20 µl. Mix and spin down.
10. Incubate the RT reaction as follows:

| **Step** | **Cycles** | **Temp (°C)** | **Time (minutes)** |
|---|---|---|---|
| Reverse Transcription | 1 | 42 | 90 |
| Denature | 1 | 85 | 5 |
| Hold | 1 | 4 | ∞ |

11. Set up Amplicon multiplex PCR in the Mastermix hood.

| **No.** | **Component** | **Volume/sample (µl)** | **5 samples (vol, µl)** |
|---|---|---|---|
| 1. | Water | 18.7 | |
| 2. | 2x LongAmp Taq Master Mix | 25 | |
| 3. | PCR primer (from SQK-PCB109)* | 1.5 | |
| 4. | cDNA | 5 | |
| | **Total Volume** | **50** | |

*Barcodes are assigned BC01, BC02, BC03, BC04, and BC05, to samples A,B,C,D, and -Ve respectively.

12. Transfer 43.5 µl of RT mix to a new tube. Add respective barcode.
13. Add 5 µl cDNA to each tube and mix well by pipetting
14. Run PCR

| Step | Cycles | Temp (°C) | Time (minutes) |
|---|---|---|---|
| Heat Activation | 1 | 95 | 30 sec |
| Denature | 20 | 95 | 15 sec |
| Annealing | | 62 | 15 sec |
| Extension | | 65 | 6 |
| Polish | 1 | 65 | 6 |
| Hold | | 4 | ∞ |

15. Add 1 µl of NEB exonuclease 1 and incubate at 37 °C for 15 minutes followed by 80 °C for 15 minutes.
16. Do cleanup using 0.8X AMPure XP beads, elute in 12 µl of EB buffer.
17. Do some QC; the D5000 assay kit as well as the Qubit

| **Code** | **I.D** | **Vol (µL)** | **Dil** | **µL Qbt** | **Conc (ng/µL)** | **Yield (ng)** | **Vol for 37.5 ng(µL)** |
|---|---|---|---|---|---|---|---|
| A | MCF7 (100 ng) | 12 | 5 | 1 | 4.94*5=25 | 300 | 1.5 |
| B | HeLa (50 ng) | 12 | 5 | 1 | 9.58*5=48 | 576 | 0.78 |
| C | GM24143 (83 ng) | 12 | 5 | 1 | 9.44*5=47 | 564 | 0.8 |
| D | Cc-E-6H (50 ng) | 12 | 5 | 1 | 8.38*5=42 | 503 | 0.89 |
| -Ve | Water | 12 | 5 | 1 | LOW | - | |

18. Pool all samples by taking 37.5 ng per sample to **a total of 150 ng** in a PCR tube. Make the volume up to 23 µL with EB.
19. Do some D5000 assay QC.
20. Add 1 µL of Rapid Adapter (RAP) to the sample and incubate for 5 minutes at RT.
21. Follow the priming and loading steps and final library preparation.

**Panhandle primer protocol: B002-05-6C2**

**For lib prep I will follow the Direct cDNA Native Barcoding (SQK-DCS109 with EXP-NBD104 and EXP-NBD114)**

22. Using the samples above.
23. Thaw the RT and PCR reagents.
24. Setup RNA-Primer annealing reaction as follows

| | | A | B | C | D | -Ve |
|---|---|---|---|---|---|---|
| | | MCF7 | HeLa | GM24143 | Cc-E-6H | Water |
| | | volume/sample (µl) | vol (µl) | vol (µl) | vol (µl) | vol (µl) |
| 1. | RNase free Water | 8.6 | 4.6 | 8.6 | 9.1 | 9.6 |
| 2. | Total RNA sample | 1 | 5 | 1 | 0.5 | 1 |
| 3. | Oligo(dT), at 10 µM | 1 | 1 | 1 | 1 | 1 |
| 4. | dNTP (10 mM) | 1 | 1 | 1 | 1 | 1 |
| | **Total Volume** | **11.6** | **11.6** | **11.6** | **11.6** | **11.6** |

25. Mix gently and spin down briefly.
26. Run the following **pre-RT mix protocol**

| Temperature (°C) | Time (min) | Purpose |
|---|---|---|
| 72 | 3 | Unfolding of RNA secondary structures, Poly-T primer binding |
| 4 | 10 | Poly-T primer binds |
| 25 | 1 | Poly-T primer binds more specifically |
| 4 | Hold | |

27. Setup Reverse Transcription (RT) mastermix as follows:

| **RT mix** | | |
|---|---|---|
| **Component** | **Volume/sample (µl)** | **5 samples** |
| RNase free Water* | 0.6 | - |
| Maxima H 5X Buffer | 4.4 | 22 |
| RNaseOUT | 1 | 5 |
| TSO (10 µM)** | 2 | 10 |
| Betaine (stock: 5M) | 2 | 10 |
| Maxima H RTase | 1 | 5 |
| **Total Volume** | **10.4** | |

*I will omit this water

28. Mix gently by flicking the tube then spin down.
29. Add the 10.4 µl of the RT mix to the 11.6 µl annealed template RNA sample.
30. Incubate the RT reaction using this custom **SSIV RT protocol:**

| Temperature | Time | Cycles | Purpose |
|---|---|---|---|
| 50°C | 10 min | 1 | RT and template-switching |
| 55°C | 30 sec | 10 | Unfolding of RNA secondary structures |
| 50°C | 30 sec | | Completion/continuation of RT |
| 60°C | 30 sec | 5 | Unfolding of RNA secondary structures |

| | | | |
|---|---|---|---|
| 55°C | 30 sec | | Completion/continuation of RT |
| 50°C | 30 sec | 1 | Finish template switching |
| 65°C | 30 sec | 5 | Unfolding of RNA secondary structures |
| 60°C | 30 sec | | Completion/continuation of RT |
| 50°C | 30 sec | 1 | Finish template switching |
| 70°C | 30 sec | 5 | Unfolding of RNA secondary structures |
| 65°C | 30 sec | | Completion/continuation of RT |
| 50°C | 30 sec | 1 | Finish template switching |
| 75°C | 30 sec | 5 | Unfolding of RNA secondary structures |
| 70°C | 30 sec | | Completion/continuation of RT |
| 50°C | 1 min | 1 | Final finish template switching |
| 80°C | 10 min | 1 | Enzyme inactivation |
| 4°C | Hold | 1 | |

31. Set up Amplicon multiplex PCR in the Mastermix hood.

| No. | Component | Volume/sample (µl) | 5 samples (vol, µl) |
|---|---|---|---|
| 1. | Water | 19 | |
| 2. | 2x LongAmp Taq Master Mix | 25 | |
| 3. | PCR primer (10 µM) | 1 | |
| 4. | cDNA | 5 | |
| | **Total Volume** | **50** | |

32. Transfer 45 µl of PCR mastermix to 3 new plates.
33. Add 5 µl of cDNA to each corresponding well and mix well by pipetting
34. Run PCR

| Temperature | Time | Cycle |
|---|---|---|
| 95°C | 1 min | 1 |
| 95°C | 20 sec | 5 |
| 58°C | 4 min | |
| 68°C | 6 min | |
| 95°C | 20 sec | 15 cycles |
| 64°C | 30 sec | |
| 68°C | 6 min | |
| 72°C | 10 min | 1 |
| 4°C | Hold | 1 |

35. Add 1 µl of NEB exonuclease 1 and incubate at 37 °C for 15 minutes followed by 80 °C for 15 minutes.
36. Do 1X AMPure. Elute in 30 µl. This is the **FIRST AMPure**.
37. Do a LabChip on all samples to see cDNA profile.
    a. Transfer 13 µl of water to corresponding wells of a 384-well plate.
    b. Transfer 2 µl (2:15 dilution) of each samples to 384-well plate.

c. Run LabChip

38. Do Qubit. Take forward 500 ng per sample for End-repair and dA tailing.

39. Do Ultra II End repair and dA tailing.

a. Prepare End repair and dA-tailing mix as follows.

**End repair and dA-tailing setup**

| No. | Component | Volume/sample (µl) |
|---|---|---|
| 1. | cDNA-PCR sample | 50 |
| 2. | Ultra II End-prep reaction buffer | 7 |
| 3. | Ultra II End-prep enzyme mix | 3 |
| | **Total Volume** | **60** |

b. Set a 100 µl or 200 µl pipette to 50 µl and then gently pipette the entire volume up and down at least 10 times to mix thoroughly. Perform a quick spin to collect all liquid from the sides of the tube.

c. Incubate as follows.

| Temperature (°C) | Time (min) | Purpose |
|---|---|---|
| **Heat lid to 75 °C** | | |
| 20 | 30 | Reaction |
| 65 | 30 | Inactivation |
| 4 | Hold | |

40. Ligate barcodes as follows.

a. Prepare Native barcode ligation mix.

**Barcoding ligation setup**

| No. | Component | Volume/sample (µl) |
|---|---|---|
| 1. | End Prep Reaction Mixture | 60 |
| 2. | Native barcode | 2.5 |
| 3. | Ultra II Ligation Master Mix | 30 |
| 4. | Ligation Enhancer | 1 |
| | **Total Volume** | **93.5** |

b. Mix the Ultra II Ligation Master Mix by pipetting up and down several times prior to adding to the reaction.

c. Add the respective barcode to each sample.

d. Add the 31 µl of ligation mix directly to each End-prep reaction.

e. Set a 100 µl or 200 µl pipette to 80 µl and then pipette the entire volume up and down at least 10 times to mix thoroughly. Perform a quick spin to collect all liquid from the sides of the tube.

f. Incubate at RT for 15 minutes then 70°C for 10 minutes

41. **Second AMPure** cleanup: 0.8X, 75 µl of AMPure, elute in 20 µl water. Quantify by Qubit.

42. Pool all samples by taking 5 µl per sample except for A where I take 4.3 µl. Make up the volume to 65 µl.

43. I skipped the **Third** round of 0.8X AMPure.

44. Do sequencing adapter ligation as follows.

a. Make the sequencing adapter ligation mix

**Sequencing adapter ligation setup**

| No. | Component | Volume/sample (µl) |
|---|---|---|
| 1. | 200 fmol pooled barcoded sample | 65 |
| 2. | Adapter Mix II (AMII) | 5 |
| 3. | NEBNext Quick Ligation Reaction Buffer (5X) | 20 |
| 4. | Quick T4 DNA Ligase | 10 |
| | **Total Volume** | **100** |

b. Incubate at RT for 15 minutes.

45. **Fourth AMPure** cleanup: 0.5X, 50 µl of AMPure, 140 µl Wash buffer x2, elute in 26 µl ElutionBuffer (EB). Quantify and profile on Tapestation.

46. Do some QC; the D5000 assay kit as well as the Qubit

| No. | Sample ID | Vol (µl) | Dil | µl Qbt | Conc'n ng/µl | Yield (ng) | Loaded (ng) | %ge recovery |
|---|---|---|---|---|---|---|---|---|
| **At step 36** | | | | | | | | |
| A | MCF7 (100 ng) | 30 | 5 | 1 | 10.6*5=53 | 1,590 | - | - |
| B | HeLa (50 ng) | 30 | 5 | 1 | 8.62*5=43 | 1,290 | - | - |
| C | GM24143 (83 ng) | 30 | 5 | 1 | 9.44*5=47 | 1,410 | - | - |
| D | Cc-E-6H (50 ng) | 30 | 5 | 1 | 6.44*5=32 | 960 | - | - |
| -Ve | Water | 30 | 5 | 1 | LOW | - | - | - |
| **At step 41** | | | | | | | | |
| A | MCF7 (100 ng) | 20 | 1 | 1 | 23.6 | 472 | 500 | 94.4 |
| B | HeLa (50 ng) | 20 | 1 | 1 | 20.2 | 404 | 500 | 80.8 |
| C | GM24143 (83 ng) | 20 | 1 | 1 | 20.0 | 400 | 500 | 80 |
| D | Cc-E-6H (50 ng) | 20 | 1 | 1 | 21.4 | 428 | 500 | 85.6 |
| -Ve | Water | 20 | 1 | 1 | LOW | - | - | - |
| **At step 46** | | | | | | | | |
| PH | B002-05-6C2 | 26 | 1 | 1 | 10.3 | 267.8 | 400 | 67 |

In both Nanopore and Panhandle protocols, I used 150 ng to load on the flow cell. Flow cell used for B002-05-6C1 had 2521 pores at beginning of sequencing. Flow cell used for B002-05-6C2 had 4593 pores at beginning.
